# Environmental contributions to the evolution of trait differences in *Geum triflorum*: implications for restoration

**DOI:** 10.1101/2022.02.11.480132

**Authors:** Kate Volk, Joseph Braasch, Marissa Ahlering, Jill A. Hamilton

## Abstract

**Premise of the Study:** Understanding how environment influences the distribution of trait variation across a species’ range has important implications for seed transfer during restoration. Heritable genetic differences associated with environment could impact fitness when transferred into new environments. Here, we test the degree to which the environment shapes the evolution and distribution of genetic effects for traits important to adaptation.

**Methods:** In a common garden experiment, we quantified trait differentiation for populations of *Geum triflorum* sourced from three distinct ecoregions and evaluated the ability of climate to predict trait variation. Populations were sourced from alvar ecoregions which experience predictable extremes in seasonal water availability and the prairie ecoregion which exhibits unpredictable changes in water availability.

**Key Results:** Plants sourced from alvar ecoregions exhibited smaller but more numerous stomata and greater intrinsic water use efficiency relative to prairie plant populations supporting the evolution of ecotypic differences. Estimates of standing genetic variance and heritable genetic variation for quantitative traits suggest alvar populations have greater adaptive potential. However, reduced evolvability suggest all populations of *G. triflorum* may have limited capacity to evolve in response to environmental change.

**Conclusions:** These results point towards the importance of understanding the role of environment in shaping the distribution and evolution of genetic differences across seed populations and how these data may inform recommendations for seed transfer across novel environments and our expectations of populations’ adaptive potential.

## INTRODUCTION

Understanding how the environment influences trait variation is essential, particularly within the context of restoration (Wang et al., 2010). The evolution of ecotypic differences for vegetative, physiological, or reproductive life history traits can lead to differential success following seed transfer across environments during restoration (McKay et al., 2005; Anderson et al., 2016; Braasch et al., 2021; VanWallendael, Lowry & Hamilton, 2022). In addition, the history of selection may influence the distribution of genetic variance of traits important to adaptation. Variance in the heritability or evolvability of traits is expected to impact the success of ecotypes when planted in novel restored environments (Broadhurst et al., 2008; Crowe & Parker, 2008; Havens et al., 2015). Thus, quantifying how environment contributes to the evolution of trait differences and the distribution of genetic variance provides important insight into contemporary adaptation and future adaptive capacity (Broadhurst et al., 2008; Bucharova et al., 2019; Hamilton et al., 2020; Kulbaba et al., 2021). This is particularly important to restoration, which aims to establish populations resilient to change.

Trait differences arise through a combination of deterministic and stochastic processes (Kawecki and Ebert, 2004; Crow et al 2018; Galliart et al., 2018). For example, climatic gradients have contributed to ecotypic differentiation among grass species for morphological (Aspinwall et al., 2013; Olsen et al., 2013), phenological (Lowry et al., 2019), physiological (Aspinwall et al., 2013; Maricle et al., 2017), and fitness traits (McMilan,1959; Galliart et al., 2018). To establish seed transfer recommendations during restoration, teasing apart the contributions of environment to the evolution of trait differences may be useful to predicting populations’ response to new environments. In this study, we focus on the evolution of physiological traits, which may evolve in response to varying water availability (Dudley et al., 1996; Picotte et al., 2007; Dittberner et al., 2019). Range wide variation in *Arabidopsis thaliana* for stomatal characteristics suggests that climatic factors have led to the evolution of changes in stomatal size and density (Dittberner et al., 2019). With smaller, but more numerous stomata, plants have a greater ability to respond rapidly to changing water availability associated with increased temperatures (Drake et al., 2013; Dittberner et al., 2019). Thus, variation in physiological traits may correspond with the evolution of ecotypes associated with environments across a species’ range.

The history of selection, particularly the degree to which environmental heterogeneity has been predictable or unpredictable across a species’ range, may impact the distribution of genetic variation underlying traits and consequently their capacity to adapt. Here, we define environmental predictability as repeatable seasonal cues associated with a given climate variable (Reed et al., 2010). Theory suggests that where populations have experienced predictable environmental cues, heritable genetic variance for phenotypic traits will increase as the total phenotypic variance is reduced (Fig. 1; Levins, 1963; Reed et al., 2010; Baythavong, 2011; Kulbaba et al., 2021). In such a scenario, heritable trait differences among ecotypes may lead to increased risk of maladaptation when seed is transferred to new environments (Reed et al., 2010; March-Salas et al., 2019). In contrast, populations sourced from unpredictable environments are expected to exhibit greater plasticity and reduced trait heritability (Fig 1; Chevin et al., 2010; Reed et al., 2010; Ghalambor et al., 2007; Baythavong, 2011; March-Salas et al., 2019). If adaptive, plasticity enables plants to modify their phenotype in response to the changed environment to maintain fitness (Reed et al., 2010; Baythavong, 2011; Becklin et al., 2016; March-Salas et al., 2019). If plasticity is non-adaptive it may come with a fitness cost (Gilbert et al., 2019). Evolvability, which is the expected change in a trait per generation for a given selection coefficient (Hansen and Houle, 2008; Hansen et al., 2011), quantifies how rapidly adaptation is predicted in a continuously shifting environment (Shaw and Etterson, 2001; Kulbaba et al., 2021). Therefore, quantifying trait heritability and evolvability for seeds sourced from predictable and unpredictable environments provides complimentary metrics to predict populations’ capacity to respond to changing selective pressures. These metrics can be used to guide seed transfer recommendations and aid in determining both the initial risk of transfer across environments and the likelihood populations will adapt once established.

**Fig 1.**
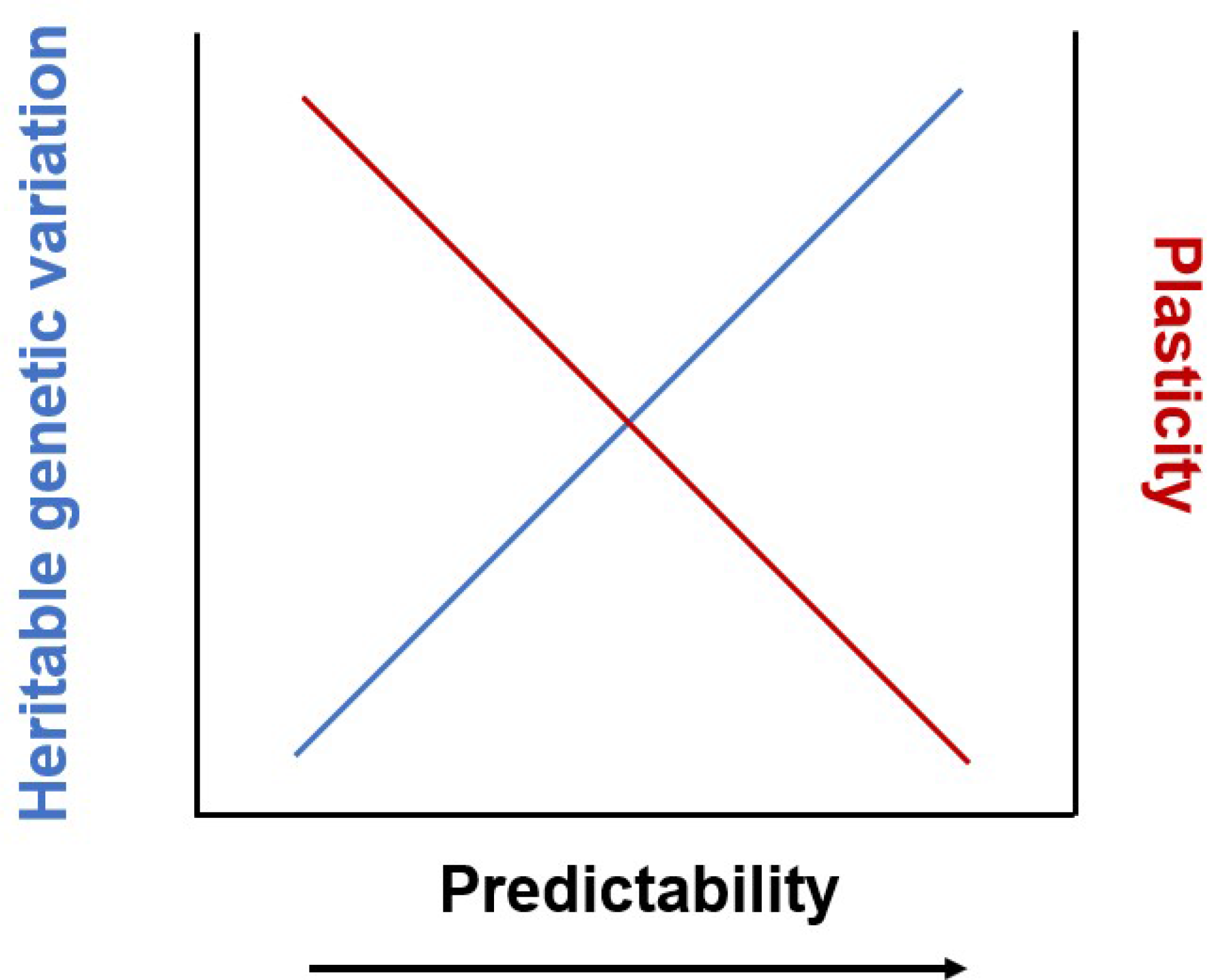
Predictive scenarios for how the distribution of heritable genetic variation could be influenced by environmental predictability. Under greater environmental predictability, the distribution of heritable genetic variation should increase (blue line) while decreasing variation attributed to plasticity (red line).

### Geum triflorum

Pursch., is an early season perennial forb associated with remnant prairie habitat across much of the Great Plains of North America (Hamilton and Eckert 2007, Yoko et al., 2020). The Great Plains are critically imperiled due to habitat loss associated with land conversion, fragmentation, and urban expansion and thus are important habitats for restoration efforts (Hoekstra et al., 2005; Gascoigne et al., 2011; Comer et al., 2018; Wimberly et al., 2018; Bengtsson et al., 2019). Populations of *G. triflorum* also persist as isolated ‘islands’ across alvar habitats scattered throughout the Great Lakes and into Manitoba, Canada (Hamilton and Eckert 2007; Yoko et al., 2020). Alvars are habitats characterized by a thin layer of soil over limestone bedrock that harbor a unique assemblage of plants largely disjunct from the core of their distribution (Hamilton and Eckert 2007). Alvars experience extreme, but predictable annual fluctuation in water availability from flooding in the spring to early summer desiccation (Catling and Brownell 1995; Hamilton et al., 2002; Yoko et al., 2020). In contrast, while prairies experience flooding and drought, compared to the predictable interannual extremes of the alvar ecoregion, the onset of these events is less predictable. In addition, the deep, organically rich soil characterizing prairie ecoregions provides a buffer to extreme water fluctuations (Anderson, 2006). Thus, we suggest the alvar ecoregion reflects a ‘predictable’ history of selection, whereas the prairie ecoregion reflects an ’unpredictable’ history of selection in response to changing water availability. These ecoregions provide an ideal system to evaluate the role predictability of the environment may play in influencing the amount and distribution of genetic variance for phenotypic traits. Physiological traits, including stomatal size and density along with water-use efficiency (WUE) are expected to vary between prairie and alvar ecoregions. Given the importance of stomatal traits and WUE to plant persistence, examining how environment of origin has influenced variation in these traits will inform seed transfer recommendations.

Using a common garden experiment of maternal seed families for *G. triflorum* sourced from both prairie and alvar ecoregions, we evaluated the role source environment has had on the distribution of physiological trait variation linked to plant water use. We quantified ecoregional differentiation for each trait and tested for correlations between functional traits and climate of origin for all sampled populations. Lastly, we quantified standing genetic variance for stomatal traits, including estimates of heritability and evolvability. Specifically, we ask 1) do physiological traits exhibit ecoregional differences, 2) is there a relationship between physiological trait variation and source climate, and 3) does the history of selection associated with seed source environment structure the distribution of additive genetic variance and the heritability or evolvability of physiological traits? We predict alvar ecoregions will exhibit smaller, but more numerous stomata in addition to greater WUE relative to prairie populations. We also expect populations from the alvar ecoregions will have greater heritability for quantitative traits relative to prairie populations due to the history of selection associated with predictable changes in water availability. An understanding of how differentiation in physiological traits evolve and the role selection may play in shaping the distribution of heritable trait variation will be valuable for predicting the response of seeds to restored environments and estimating their longer-term evolutionary potential.

## MATERIALS AND METHODS

### Field sampling

In the spring of 2015, seed from 19 populations of *Geum triflorum* Pursch. (Rosaceae) spanning much of its distribution were collected, including 11 populations from the Great Lake alvar ecoregion (GLA), two from the Manitoba alvar ecoregion (MBA), and six from the midwestern prairie ecoregion (PRA, Fig. 2). Forty individual seed heads, each representing a maternal family, were harvested approximately every two meters along a 100 m transect within each population (as in Hamilton and Eckert 2007). In addition to field collections, three seed populations with known provenance were provided by commercial growers from within the prairie ecoregion and incorporated. Two populations were provided by commercial growers (SD- PMG, MN-PMG), and one from the United States Department of Agriculture collected near Pullman, WA (WA-BLK). In total, twenty-two populations were sampled across much of the species’ distribution for inclusion in the common garden experiment (Fig. 2).

**Fig 2.**
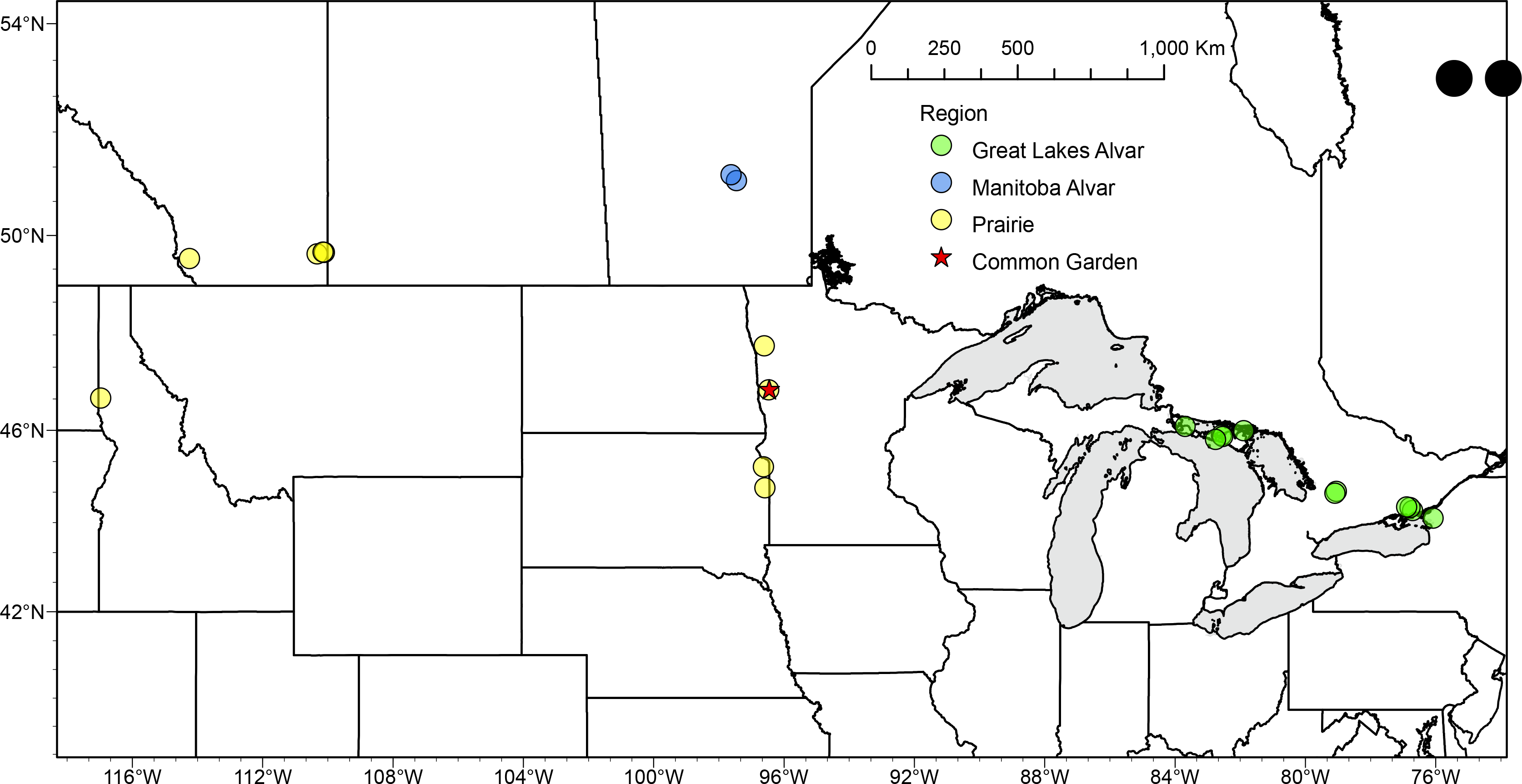
Collection sites of *G. triflorum* populations. Green points represent Great Lake Alvar (GLA) populations, blue points represent Manitoba Alvar populations (MBA), and yellow points represent Prairie populations (PRA). The red star shape indicates the common garden location.

### Common garden experiment

On November 7^th^, 2015, seeds were planted in a greenhouse at North Dakota State University in Fargo, ND. Using a half-sib design, 12 seeds from each of ten maternal seed families from each population were planted across 12 randomized complete blocks. In addition, two individual seeds from each commercial collection were planted in each block, for a total of 24 seeds per commercial seed population. In May 2016, surviving seedlings (58% of the total, Yoko et al., 2020) were transferred to a permanent field common garden location at Minnesota State University at Moorhead’s Ecoregional Science Center (46.86913N, - 96.4522W) in Moorhead, MN. The randomized-complete block design was maintained, and a full description of the establishment and maintenance of the common garden experiment can be found in Yoko (2020).

### Measurement of physiological traits

#### Stomatal Traits

In July 2016, using mature plants from the field common garden experiment, both abaxial (lower) and adaxial (upper) leaf surfaces were assessed for stomatal trait variation. Using a thin layer of Newskin “liquid bandage,” two randomly selected leaves per individual were sampled to quantify lower and upper leaf surface stomatal trait variation (N=650, 417 GLA, 91 MBA, 142 PRA). ’Liquid bandage’ leaf impressions were mounted onto slides and photographed using a Zeiss Stereo Discovery (V8) digital microscope (Carl Zeiss Microscopy, LLC, Thornwood, NY, USA) with a Canon Rebel T3 E0S 1100D digital camera (Canon Virginia Inc., Newport News, VA, USA). Photographs were standardized to a 0.32 x 0.42 mm grid and were subsequently analyzed for stomatal trait variation using ImageJ software (v1.52a, National guar m), which is a proxy for stomatal size, stomatal density (SD, mm^2^), which represents the number of stomata per unit leaf area, and area of the leaf occupied by stomata, stomatal area index (SAI, mm^2^). Measurements are reported independently for both abaxial (ab) and adaxial (ad) leaf surfaces. Individual stomatal size measurements represent an average of three guard cells per individual. Stomatal density was calculated by dividing the total number of stomata per slide by the area of the grid. The stomatal area index is reported as the product of average guard cell length and stomatal density (Bertel et al., 2017).

#### Water-use efficiency

In May 2018, leaf samples were harvested to estimate intrinsic water-use efficiency (WUE). Foliar carbon isotopes were analyzed because they provide an ability to assess water-use efficiency over the lifetime of a leaf (Farquhar et al., 1989). We sampled leaves from approximately five individuals per population (53 GLA, 9 MBA and 31 PRA). Leaf tissue was oven-dried at 55LJ over 24 hours and then homogenized into a fine powder using a TissueLyser II (Qiagen, Hilden, Germany). Between 4-5 mg of homogenized leaf tissue were weighed and placed into a tin capsule (Costech, Valencia, CA, USA) for ^13^C isotope analysis using a continuous-flow isotope ratio mass spectrometer (Sercon Ltd., Cheshire, UK) at UC Davis Stable Isotope Facility (Davis, CA, USA). The reported Pee Dee Belemnite.

### Statistical analysis

C values are expressed relative to the Vienna

### Assessing genetic differences across ecoregions

To test for differentiation in stomatal traits and WUE associated with ecoregion of origin, we used an analysis of variance (ANOVA). All traits were first assessed for normality and homogeneity of variance. Following the ANOVA, a posthoc Tukey honest significance test was performed to identify significant pairwise differences between ecoregions. All statistical tests were performed in R (R Core Team, 2018).

### The role of climate to population trait variation

To evaluate the relationship between climate of origin and its contribution to physiological trait variation, we extracted annual climatic variables, representing the most recent 30-year averages spanning 1980-2010, from ClimateNA v.5.50 using source population latitude, longitude, and elevation (Wang et al., 2016). Climate variables were highly correlated; therefore, we performed a principal component analysis (PCA) to reduce multicollinearity across traits using R (R Core Team, 2018). PC1 explained 47% of the climatic variation across populations, and PC2 explained 30%. Given the first two PC axes explained over 77% of the variation across traits, subsequent analyses reflect only PC1 and PC2. (Supporting Table 1).

**Table 1.** Summary of ANOVA for stomatal traits and WUE comparing differences associated with ecoregion of origin for Geum triflorum seedlings planted in a common environment. Bold values are significant P<0.001

| **Trait** | **df** | **SS** | **F** | **P** |
| --- | --- | --- | --- | --- |
| GCLab | 2.00 | 1.5E+02 | 19.15 | **<0.001** |
| GCLad | 2.00 | 2.7E+02 | 29.83 | **<0.001** |
| SDab | 2.00 | 2.7E+05 | 29.12 | **<0.001** |
| SDad | 2.00 | 1.6E+05 | 35.29 | **<0.001** |
| SAIab | 2.00 | 8.7E+01 | 27.59 | **<0.001** |
| SAIad | 2.00 | 6.7E+01 | 24.75 | **<0.001** |
| WUE | 2.00 | 9.1E+00 | 12.12 | **<0.001** |

Note: GCLab, abaxial guard cell length; GCLad, adaxial guard cell length; SDab, abaxial stomatal density; SDad, adaxial stomatal density; SAIab, stomatal area index; WUE, water-use efficiency

To test whether differences in stomatal traits and WUE could be explained by climatic variation summarized as PC1 and PC2, we fit linear regressions using the lm function in R (R Core Team, 2018). We tested three models using PC1 and PC2 as predictor variables. Two models included each PC as a single predictor, and the third model included both factors as additive predictors. The best-fitting model was identified as that with the lowest Akaike Information Criterion (AIC, Supporting Table 1).

### Ecoregion-specific genetic variance for stomatal traits

Additive genetic variance (V_A_), broad-sense heritability (*H^2^),* narrow-sense heritability (*h^2^),* and evolvability (CV_A_) were estimated for all stomatal traits. As only a subset of individuals were assessed for WUE, it was not possible to estimate genetic variance components for this trait. Phenotypic variance attributable to ecoregion, population, family, and block were evaluated using a linear mixed-effect model and the lmer_alt function within the afex package in R (Singmann et al., 2021):

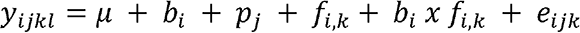

where is the phenotypic variance attributable to the random effect of the *i*th block, is the random effect of the *j*th population, is the random effect of the *k*th family within the *i*th population, is the interaction effect of block x family, and is the random error. We calculated the additive genetic variance (V_A_) as:

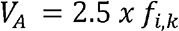

Where, is the random effect of the *k*th family within the *i*th population. Traditionally, narrow-sense heritabilities use a coefficient of relationship (p=1/4) to account for a half-sib design like that used in our experiment. However, we modified our model based on Ahrens et al., (2020) to account for the potential mixed mating system of *G. triflorum* adopting a coefficient of relationship of p=1/2.5 to generate conservative values.

Broad-sense heritability (*H*^2^) was calculated as follows using variance components from the mixed model:

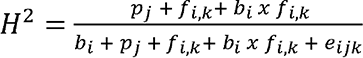

The variance components associated with the *k*th family within the *i*th population and the block x family (*b*_*i*_ *x f*_i,k_) interaction effect was subsequently used to estimate narrow-sense heritability and the evolvability. Narrow-sense heritability (*h^2^*) for leaf surface traits was estimated using the following equation:

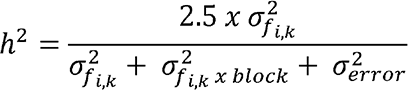

where *h^2^* is the narrow-sense heritability 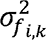, is the family within population component of 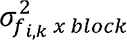 is the variance in the interaction effect of block and family (*b_i_, x f_i,k_*) and is the error component (*e_ijk_*) representing individual variance. Standard errors for *h* were calculated by dividing the pooled standard deviations of the model by the trait sample size.

In addition to broad and narrow-sense heritability estimates, we included estimates of evolvability to standardize comparisons of additive genetic variance as a means for comparison across traits and ecoregions (Hansen and Houle 2008; Hansen et al., 2011; Cava et al., 2019). We calculated evolvability as follows:

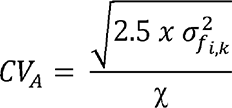

Where 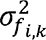 is the family component of variance, and χ is the mean of the trait under evaluation.

## RESULTS

### Physiological trait differentiation

Using an ANOVA, we tested for differences in stomatal traits and WUE for populations of *G. triflorum* sourced from distinct ecoregions. Significant trait differentiation was observed between ecoregions for all traits (Table 1). Populations from both GLA and MBA ecoregions exhibited reduced guard cell length for abaxial (GLA=27.1 µm± 0.1 µm; MBA=26.5 µm ± 0.2 µm) and adaxial (GLA=26.5 µm ± 0.1 µm; MBA=26.1 µm ± 0.2 µm) leaf surfaces relative to PRA values (abaxial 28.2 µm ± 0.2 µm and adaxial, 28.1 µm ± 0.2 µm, Fig 3A-B). While populations sourced from both GLA and MBA did not differ significantly in abaxial stomatal density, both had greater overall stomatal density (260.8 mm^2^ ± 3.7 mm^2^; 246.3 mm^2^ ±7.5 mm^2^ respectively) relative to populations sourced from the PRA ecoregion (PRA=206.6 mm^2^ ±5.8 mm^2^, Fig3C). Interestingly, all three ecoregions differed significantly from each other for adaxial stomatal density, with MBA exhibiting the greatest (MBA=193.3 mm^2^ ± 5.3 mm^2^), followed by GLA (GLA=176.6 mm^2^ ± 2.6 mm^2^), and PRA (PRA=141.32 mm^2^ ± 4.1 mm^2^, Fig3D). For stomatal area index (SAI) significant ecoregional differences were observed across both leaf surfaces (Fig3E-F). Populations sourced from the GLA ecoregion had, on average, the largest abaxial stomatal area index (GLA=5.28 mm^2^ ± 01 mm^2^), followed by MBA (MBA=4.87 mm^2^ ± 0.1 mm^2^), with PRA exhibiting the lowest (PRA=4.30 mm^2^ ±0.1 mm^2^). For adaxial stomatal area index, MBA exhibited the greatest SAI (MBA=5 mm^2^± 0.1 mm^2^), followed by GLA (GLA=4.7 mm^2^ ±0.1 mm^2^), while PRA consistently exhibited reduced SAI for both leaf surfaces relative to alvar ecoregions (PRA=3.93 mm^2^ ±0.1 mm^2^). Plants sourced from GLA exhibited higher water use efficiency (GLA=-29.4 ± 0.1) relative to MBA (-29.7 ±0.2), although there was no significant difference between the two alvar ecoregions. In contrast, plants sourced from the PRA ecoregion exhibited significantly reduced water use efficiency relative to both GLA and MBA (-30.1 ±0.1) (Fig4).

**Fig 3.**
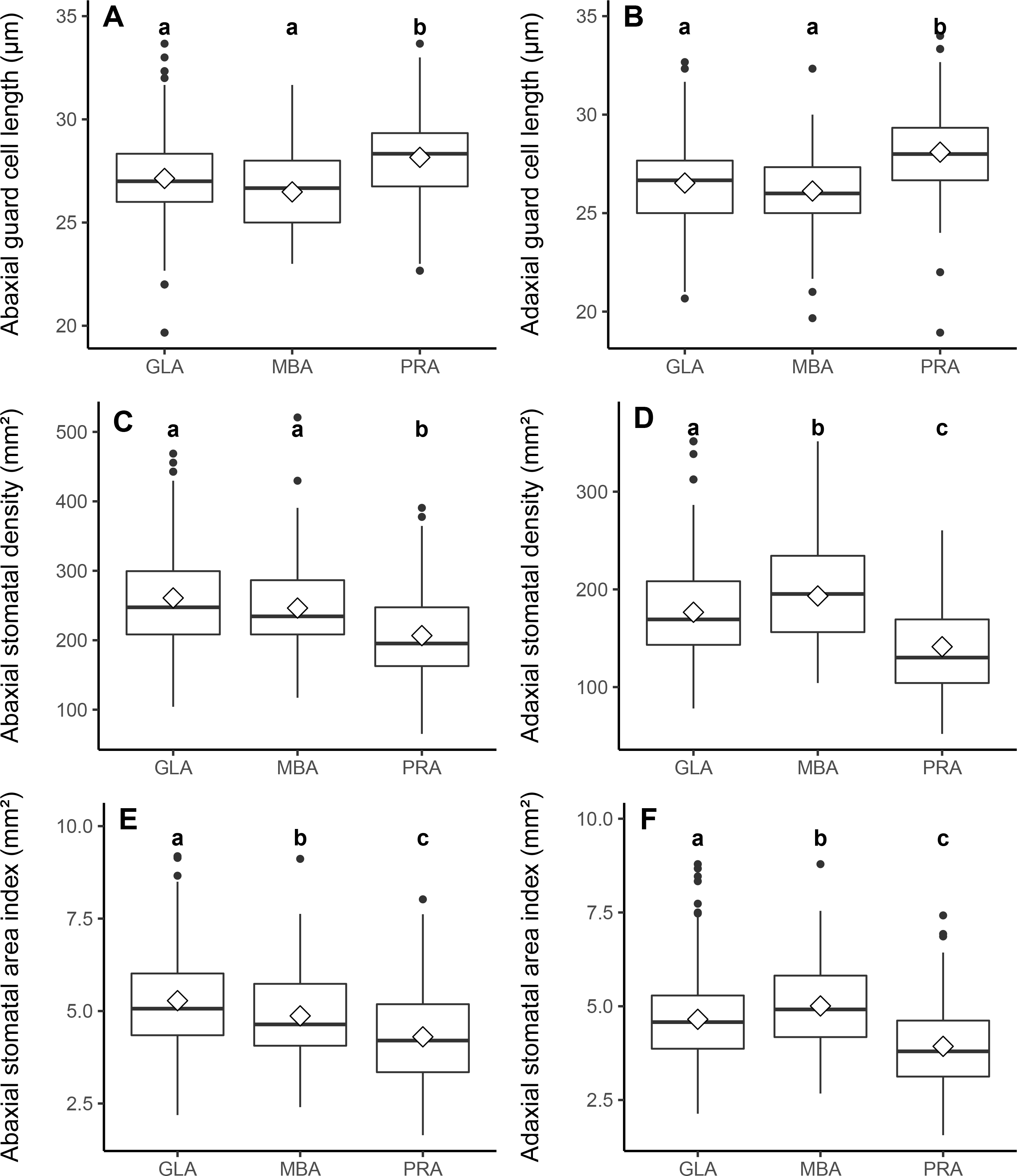
Box plots indicate regional differences in physiological traits associated with water-use, including abaxial guard cell length (A), adaxial guard cell length (B), abaxial stomatal density (C), adaxial stomatal density (D), abaxial stomatal area index (E), and adaxial stomatal area index (F). The horizontal line in the box plot indicates the median, and white diamonds indicate the mean. Boxplots with the same letter are not significantly different based on Tukey’s comparison of means (alpha = 0.05).

**Fig 4.**
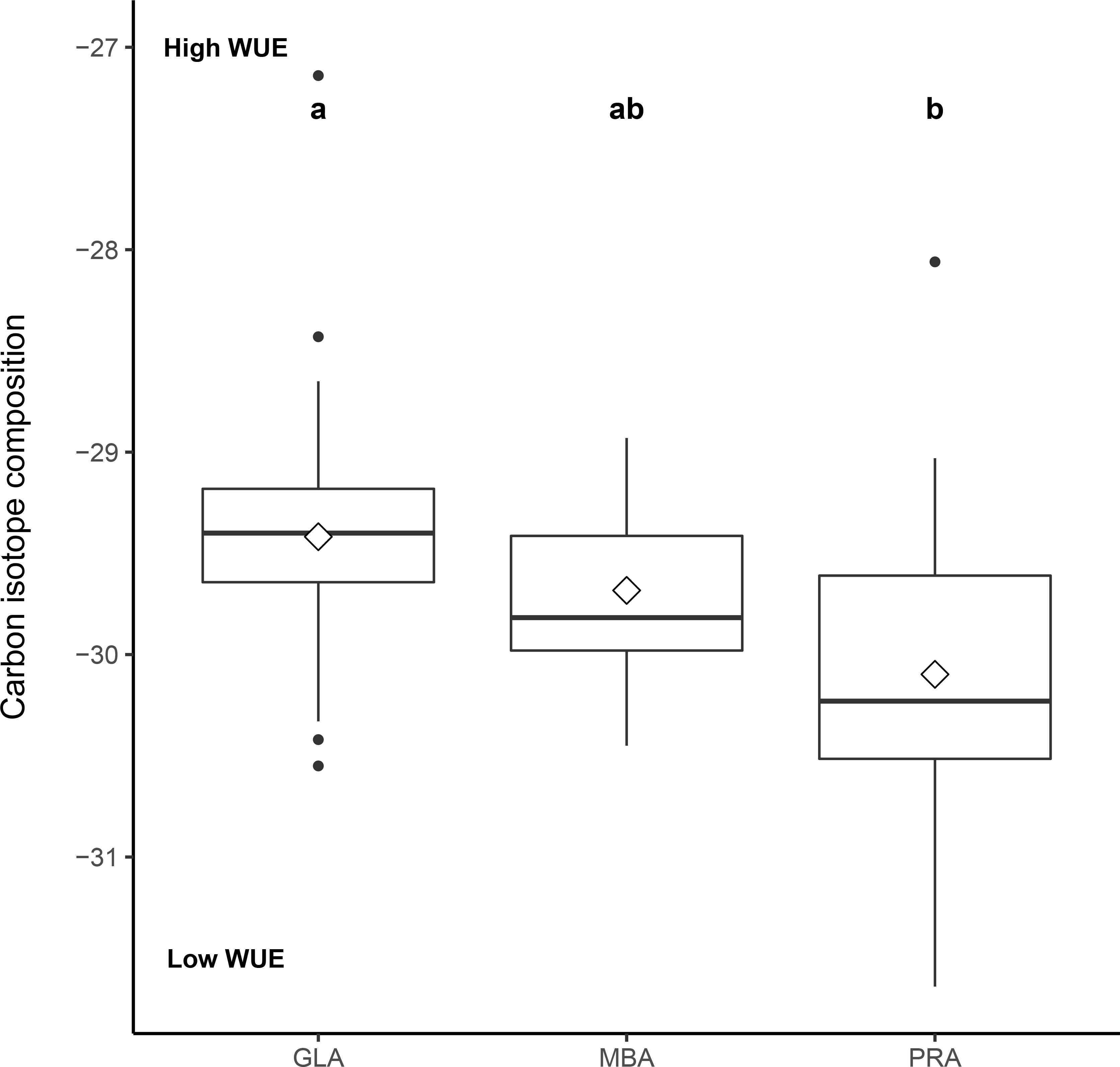
Box plot indicates regional differences in water-use efficiency (WUE). The horizontal line in the box plot indicates the median, and white diamonds indicate the mean. Boxplots with the same letter are not significantly different based on Tukey’s comparison of means (alpha = 0.05).

### Physiological trait variation associated with climate of origin

Using a PCA based on 30-year climate averages we assessed the distribution of climate variation for populations sourced from prairie and alvar ecoregions (Table S1). The first two axes of the PCA explained over 77% of the variation, with PC1 explaining 47% and PC2 explaining 30%. The climatic variables that exhibited the highest loadings on PC1 were all associated with temperature and photoperiod, including mean annual temperature (MAT), growing degree days above 18°C (DD_18), and variables related to the frost-free period (eFFP and FFP). Climate variables with the highest loadings on PC2 were related to water availability, including climate-moisture deficit (CMD) and annual heat moisture index (AHM) which indicate the amount of water available for plant uptake. A linear regression with combinations of PC1 and PC2 was used to predict stomatal trait and WUE variation for plants grown in the common garden. The model that best predicted physiological trait variation was assessed by comparing AIC scores, with the chosen model exhibiting the lowest AIC value. In nearly all cases PC2 was the best predictor for stomatal traits and WUE (Table S2), indicating water availability likely explains variance in physiological traits for *G. triflorum.* Populations sourced from regions of the species’ range that experience increased climate moisture deficit and higher annual heat moisture indices were associated with increased abaxial and adaxial guard cell length (r^2^=0.39, *P*<0.01; r^2^=0.39, *P*<0.01 respectively, Fig5 A-B). In contrast, greater climate moisture deficit and higher annual heat moisture associated with the PC2 axis predicted reduced stomatal density (r^2^=0.40, *P*<0.01; r^2^=0.37, *P*<0.01 respectively, Fig5 C-D). This indicates that populations from ecoregions with reduced water availability on average produced fewer, but larger, stomata.

**Table 2.** Ecoregional-specific broad-sense heritabilities (H^2^), additive genetic variance (V_A_), narrow-sense heritabilities (h^2^) and evolvability (CV_A_)

| Trait | PRA |  |  |  | MBA |  |  |  | GLA |  |  |
| --- | --- | --- | --- | --- | --- | --- | --- | --- | --- | --- | --- |
| | $H^2$ | $V_A$ | $h^2$ | $CV_A$ | $H^2$ | $V_A$ | $h^2$ | $CV_A$ | $H^2$ | $V_A$ | |
| GCLab | 0.09±0.05 | 0.42 | 0.22±0.15 | 0.04 | 0.8±0.2 | 0.51 | 0.39±0.20 | 0.04 | 0.9±0.28 | 0.60 | 0 |
| GCLad | 0.12±0.1 | 0.42 | 0.21± 0.15 | 0.04 | 0.8±0.21 | 0.00 | 0.0±0.0 | 0.00 | 0.9±0.32 | 0.50 | 0 |
| SDab | 0.32±4.35 | 168 | 0.13±2.93 | 0.10 | 0.9±9.1 | 239.00 | 0.13±4.19 | 0.10 | 0.86±10.12 | 425.00 | 0 |
| SDad | 0.4±4.17 | 240 | 0.37±3.49 | 0.17 | 0.9±5.8 | 252.00 | 0.27±4.35 | 0.13 | 0.85±7.1 | 353.00 | 0 |
| SAIab | 0.34±0.09 | 0.07 | 0.17±0.06 | 0.10 | 0.9±0.12 | 0.00 | 0 | 0.00 | 0.86±0.18 | 0.10 | ( |
| SAIad | 0.33±0.12 | 0.2 | 0.44±0.10 | 0.18 | 0.84±9,15 | 0.30 | 0.52±0.15 | 0.17 | 0.85±0.16 | 0.19 | 0 |

Note: GCLab, abaxial guard cell length; GCLad, adaxial guard cell length;SDab,abaxial stomatal density; SDad, adaxial stomatal density; SAIab,stomatal areaindex; WUE, water-use efficiency; PRA, prairie; MBA, Manitic alvar

Similarly, both abaxial and adaxial stomatal area indices decreased with larger values of PC2 (r^2^=0.41, *P*<0.01; r^2^=0.33, *P*<0.01, respectively, Fig5 E-F). Finally, WUE decreased across the PC2 axis (r^2^=0.37, *P*<0.01, Fig. 6), indicating the control over water-use decreases under increased CMD and AHM for *G. triflorum* populations.

**Fig 5.**
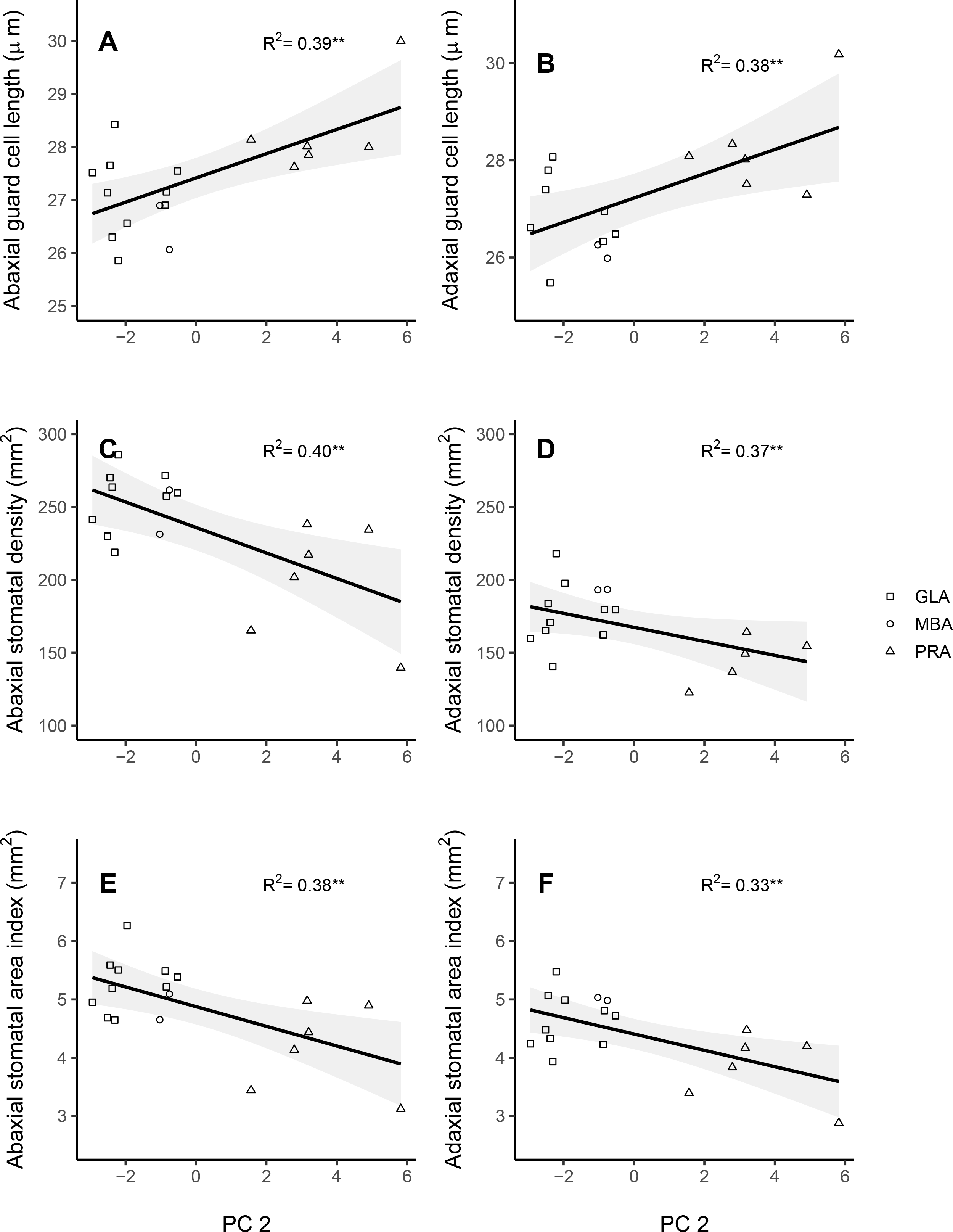
Relationships between stomatal traits and principal component 2 (PC2), including abaxial guard cell length (A), adaxial guard cell length (B), abaxial stomatal density (C), adaxial stomatal density (D), abaxial stomatal area index (E), and adaxial stomatal area index (F). Data points represent population-level averages for each region. Lines depict the shape of the association between PC2 and trait values surrounded by a 95% confidence shading. The significance of each relationship is indicated in the top right corner of each graph (p<0.01).

**Fig 6.**
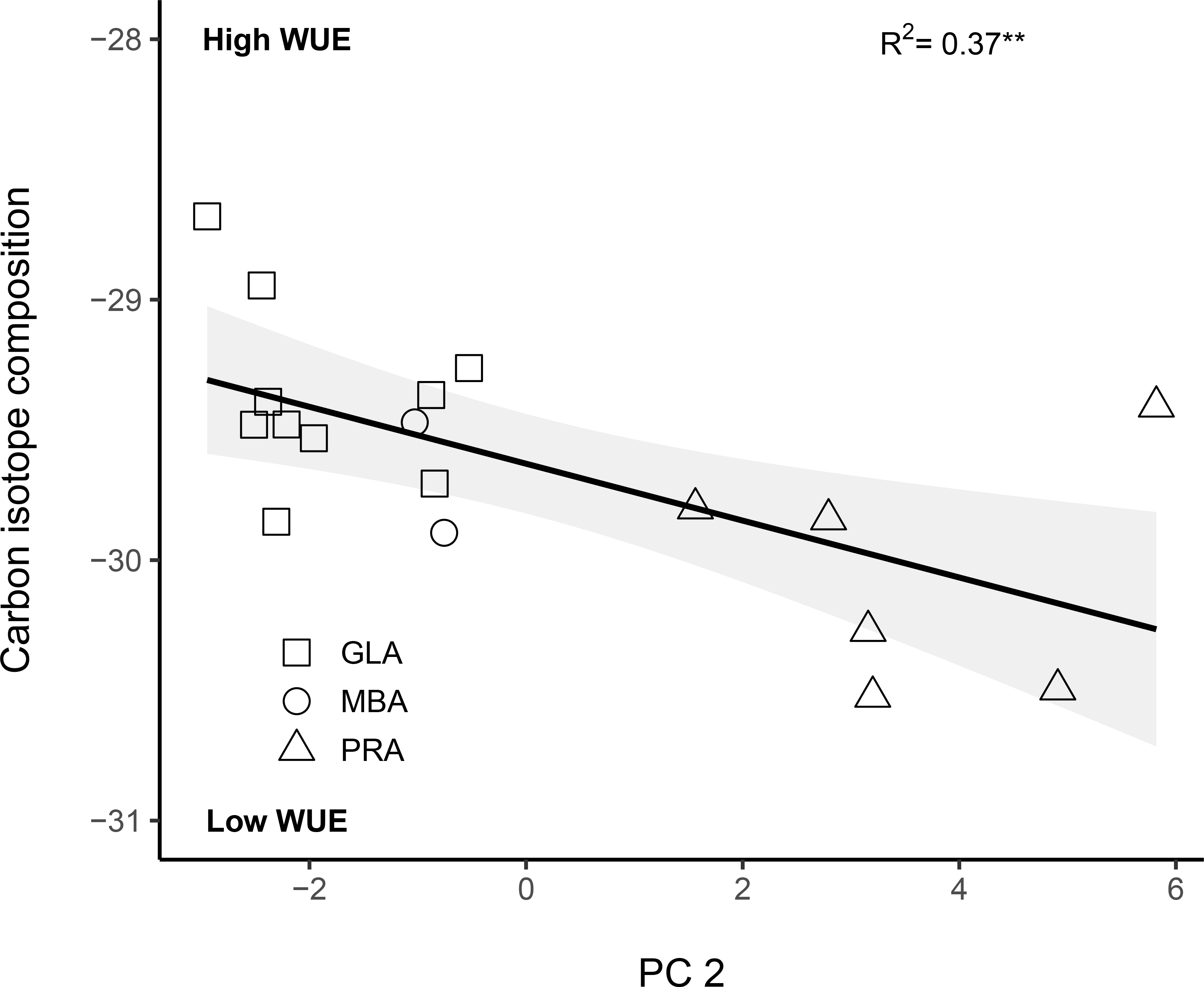
Relationships between water-use efficiency (WUE) and principal component 2 (PC2). Data points represent population-level averages for each region. Lines depict the shape of the association between PC2 and trait values surrounded by a 95% confidence shading. The significance of the relationship is indicated in the top right corner of each graph (p<0.01).

### Distribution of additive genetic variance for stomatal traits

Additive genetic variance (V_A_) provides an estimate of the amount of genetic variation available for selection to act upon (Falconer and Mackay, 1996). Ecoregion-specific estimates for additive genetic variance (V_A_) of stomatal traits were quantified using the half-sibling design (Table 2). In the common garden, Great Lake alvar individuals exhibited the greatest V_A_ for adaxial and abaxial guard cell length (0.60, 0.50), stomatal density (425, 353) and for abaxial stomatal density (0.10). These results suggest that for *G. triflorum*, the majority of standing genetic variation for stomatal traits is associated with the Great Lake alvar ecoregion.

### Heritability of stomatal traits

We estimated both broad-sense and narrow-sense heritabilities for traits across eco- regions to understand how history of selection may influence the distribution of phenotypic trait variance. Broad-sense heritability (*H*^2^) accounts for all genetic components of total phenotypic variance and was calculated for each ecoregion. Estimates of stomatal trait *H*^2^ ranged from 0.09-0.9 (Table 2). *H*^2^ was comparable for GLA and MBA individuals for abaxial and adaxial GCL (GLA= 0.9±0.28, MBA= 0.8±0.2; GLA= 0.9±0.32, MBA= 0.8±0.21, respectively), and PRA exhibited the lowest *H*^2^ for abaxial and adaxial GCL (0.09±0.05; 0.12±0.1, respectively). Across all three ecoregions, large variances were observed around *H*^2^ estimates for abaxial and adaxial stomatal density traits limiting our ability for comparison of stomatal density *H*^2^ across ecoregions. Finally, abaxial and adaxial stomatal area index showed similar trends to *H*^2^ estimates for GCL, where individuals sourced from the alvar ecoregions had comparable *H*^2^ estimates (GLA= 0.86±0.18, MBA= 0.9±0.12; GLA= 0.85±0.16, MBA= 0.84±9.15, respectively) and were greater than individuals sourced from the PRA ecoregion (0.34±0.09, 0.33±0.12). These results suggest physiological trait differences associated with ecoregions are likely attributable to genetic effects.

While broad sense heritabilities includes total genetic variance and provide an understanding of total genetic effects contributing to a trait phenotype, narrow-sense heritabilities (*h^2^*) account for the proportion of genetic variance attributed to additive effects, providing an understanding of potential response to selection. There was substantial variability in narrow sense heritabilities estimated for physiological traits across ecoregions (Table 2). On average, individuals from the GLA ecoregion exhibited greater *h^2^* for all traits, excluding stomatal area index, which was greatest for individuals sourced from the PRA ecoregion (Table 2). Narrow-sense heritability was similar between GLA (0.43±0.10) and MBA (0.39±0.20) for abaxial guard cell length and *h^2^* for individuals sourced from the GLA ecoregion were relatively consistent across leaf surfaces (adaxial GLA=0.39±0.20). However, for individuals sourced from the MBA ecoregion there was no heritability observed for adaxial guard cell length, likely reflecting a lack of maternal families for this ecoregion. For abaxial guard cell length, GLA exhibited the highest *h^2^* (0.43±0.10), followed by MBA (0.39±0.20) and PRA, which exhibited the lowest degree of *h^2^* (0.22±0.15). Additionally, GLA exhibited the greatest *h^2^* for adaxial guard cell length traits (0.39±0.10), followed by PRA (0.21± 0.15). Across all three ecoregions, large variances were observed for heritabilities for abaxial and adaxial stomatal density, hindering our ability to make ecoregion comparisons for stomatal density. For abaxial stomatal area index, PRA exhibited the greatest *h^2^* (0.17±0.06), followed by GLA (0.1±0.04), however, no *h^2^* was observed for MBA individuals as seen with adaxial guard cell length. Lastly, MBA had the greatest *h^2^* for adaxial stomatal area index (0.52±0.15), followed by PRA (0.44±0.10), and GLA exhibited the lowest (0.41±0.06). These values suggest a proportion of the ecotypic differences observed between regions are likely attributable to additive genetic effects with some variance across ecoregions.

### Evolvability of stomatal traits

To determine whether the adaptive capacity of stomatal traits differs across ecoregions we calculated evolvability. While alvar and prairie ecoregions exhibited eco-region differences in V_A_, evolvability did not vary by ecoregion (Table 2). This suggests, that while the amount of additive genetic variance is greater for alvar ecoregions, the per-generation change expected due to any given selection coefficient is similar across ecoregions. Although evolvability did not vary by ecoregion it did vary across traits (Table 2). Abaxial and adaxial guard cell length had the lowest evolvabilities (0-0.04), and adaxial stomatal area index exhibited the greatest (0.15-0.18), suggesting that the per generation change will be greater for stomatal area index traits than for guard cell length. Overall, evolvabilities for all traits were low ranging from 0- 0.18 indicating that the expected per generation change in these traits is likely limited (Table 2).

## DISCUSSION

Selection associated with environment can influence the distribution of genetic variance underlying traits across a species’ range. Here, by examining populations of *G. triflorum* sourced from distinct ecoregions with contrasting predictability in water availability, we observed substantial differentiation in physiological traits that could impact recommendations for seed transfer across environments. Climate factors associated with varying water availability strongly predicted physiological trait variation across ecoregions, indicating that environment has likely contributed to the evolution of trait differences. Populations sourced from the alvar ecoregion exhibited increased stomatal density but reduced stomatal size and greater water use efficiency relative to prairie populations when grown in a common environment. This suggests that plants sourced from alvar ecoregions may have evolved increased control over water use. In addition, additive genetic variance for physiological traits was greater for populations sourced from the environmentally predictable alvar ecoregions relative to those sourced from the prairie ecoregion. Heritability estimates suggest the alvar populations exhibit increased genetic control over the phenotypic expression of physiological traits. However, estimates of evolvability suggest that exposure to varying selection coefficients may lead to limited change in traits over generations across ecoregions. Thus, our results suggest that while the environment contributes to the evolution of genetic differences across ecoregions and the distribution of genetic variance in traits important to adaptation, the adaptive capacity overall of *G. triflorum* may be limited range wide. Combined, the evolution of genetic differences may lead to environment-trait mismatches following movement of seed across ecoregions and populations may have limited capacity to buffer the fitness consequences of mismatches via plasticity.

### Physiological trait differentiation associated with seed source environment

Physiological traits often exhibit differentiation associated with environment of origin (Dudley, 1996; Didiano et al., 2016; Dittberner et al., 2019; Galliart et al., 2018; Ramirez-Valiente et al., 2018). Here we observed that when grown in a common environment, populations sourced from the alvar ecoregion exhibited, on average, smaller and more numerous stomata relative to populations sourced from prairie ecoregions. In addition, alvar populations exhibited greater intrinsic WUE relative to prairie populations suggesting that physiological traits are differentiated across ecoregions. Alvar environments exhibit annual cycles of extreme variation in water availability; from flooding in the spring to early summer desiccation (Catling and Brownell, 1995, Yoko et al., 2020;). Thus, variation across ecoregions may reflect the evolution of physiological traits required to maintain fitness under seasonal extremes in water availability. Many, but small stomata may enable plants to respond rapidly to varying extremes within the alvar ecoregion (Drake et al., 2013). Previous studies have shown the evolution of traits in response to water stress (Anderson et al., 2011; Wadgymar et al., 2016) or have directly linked physiological trait variation and water-availability to source environment (Dudley, 1996; Didiano et al., 2016; Dittberner et al., 2019; Galliart et al., 2018; Ramirez-Valiente et al., 2018). Our results demonstrate that the environment of seed source may contribute to the evolution of phenotypic differences in physiological traits, which could impact fitness if seed is transferred to a new environment during restoration.

Stochastic evolutionary processes may also contribute to trait differentiation observed for populations of *G. triflorum*. Previous studies suggest that alvar populations were likely founded from an expansion of the prairie ecoregion during the warming Hypsithermal but have subsequently become isolated following the consequent cooling period (Hamilton and Eckert, 2007). Thus, genetic differences may have accumulated across alvar populations due to stochastic processes associated with isolation and reduced connectivity relative to prairie populations (Lande 1992, Young et al., 1996). If lack of gene flow or drift following isolation were the primary mechanisms contributing to differentiation, we would expect geographically proximal MBA populations to be more similar to PRA populations, where there is a common history and high probability of gene flow between ecoregions that would limit the evolution of trait differences. However, our results suggest that Great Lake and Manitoba alvar populations are more similar to each other, suggesting that selection associated with environment has likely driven the evolution of physiological trait differences among ecoregions.

### Climate of origin predicts physiological trait variation

Using climate associated with population origin we performed a PCA to identify those climate variables that structure population variation across the range of *G. triflorum.* We found ecoregions were differentiated primarily by temperature (PC1) and water availability (PC2).

While PC1 explained the most variation between ecoregions, it did not predict physiological trait variation. However, we did observe a relationship between PC2 and physiological trait variation. Using the PC2 axis, we observed greater annual climate moisture deficit (CMD) and annual heat moisture index (AHM) were associated with fewer, but larger stomata and lower WUE characteristic of the prairie ecoregion. Previous studies found similar patterns where drier conditions led to reduced stomatal control impacting plant water use (Didiano et al., 2016; Guo et al., 2017). Yoko (2020) suggested that stomatal traits and WUE in *G. triflorum* were likely under strong divergent selection due to ecoregional differences in water availability.

Interestingly, other traits examined by Yoko (2020) may be related to climatic variables along the PC1 axis. For example, Yoko (2020) found that prairie populations invest more energy towards resource allocation than alvar populations, which may be related to variables observed along the PC1 axis.

### Distribution of genetic variance across ecoregions

The amount of additive genetic variance in fitness-related traits is proportional to the amount of genetic variance available for selection to act upon (Kulbaba et al., 2019). Here, we found that individuals sourced from the alvar ecoregions, which exhibit predictable seasonal extremes in water availability, exhibited the greatest amount of additive genetic variance for stomatal traits (Table 2). Temporally varying, but predictable environments like those featured in the alvar ecoregion likely favor the maintenance of additive genetic variance (Levins, 1963; Baythavong, 2011; Kulbaba et al., 2021). As such, estimates of V_A_ for prairie populations, which experience unpredictable changes in water availability, may reflect selection for plasticity (Baythavong, 2011; Kulbaba et al., 2021). Increased estimates of V_A_ from the alvar ecoregion for stomatal traits also suggest that these populations may harbor greater capacity to respond to selection (Kulbaba et al., 2019). However, it is important to note the variance in V_A_ estimates across ecoregions may reflect variance in the number of families included for each regional estimate of trait variance in our common garden experiment (GLA=60; MBA=12; PRA=18).

Fewer maternal families evaluated in the MBA and PRA region may lead to conservative estimates of V_A._

### Heritability of stomatal traits

Quantifying heritability provides insight into the degree to which trait variation is largely mediated by genetic or environmental effects. We predicted that heritable trait variation would be greater within alvar environments as they experience environmental heterogeneity that is predictable. Interestingly, broad-sense heritabilities for stomatal traits ranged from 0.09-0.9 and narrow-sense heritability estimates ranged from 0.1-0.5 and were similar across ecoregions (Table 2). This suggests that while the genetic effects for some traits is substantial, there is also a substantial proportion of variance attributable to environmental variance. Indeed, unpredictable, heterogeneous environments may select for the maintenance of plasticity to ensure plant resilience to change (Chevin et al., 2010; Reed et al., 2010; Ghalambor et al., 2007; Baythavong, 2011; March-Salas et al., 2019). As maternal effects are strongest in first generation seedlings and the first year of growth (Donohue, 2009), it is possible that heritability estimates in our study would decrease in a second generation. However, the perennial life-history and time to produce seed did not facilitate the inclusion of a second generation. To limit the potential effect of the maternal environment we evaluated trait variation following at least six months of establishment in the field as previous studies have indicated the impact of the maternal environment may diminish over time (Donohue, 2009). However, given these caveats, our estimates of both broad and narrow-sense heritability likely represent upper bounds (Falconer and Mackay, 1996).

Understanding the heritability of traits remains important in the context of restoration, where the degree to which traits are mediated by genetic or environmental effects can be used to inform seed transfer guidelines (Broadhurst et al., 2008; Espeland et al., 2017; Bucharova et al., 2017). Here, we observed that individuals sourced from the alvar ecoregion exhibited greater broad-sense heritability relative to individuals from the prairie ecoregion. This may suggest that ecotypic differences that may have arisen in one environment may impact expression of phenotypes in novel restored environments. Where there is a difference between seed source and transferred environment an increased probability of environmental mismatch may reduce fitness in the restored environment (Reed et al., 2010; March-Salas et al., 2019). Despite this, narrow-sense heritability estimates indicate that populations may be able to produce a plastic response to the environment, potentially mitigating negative effects associated with seed transfer and climate change (Arntx and Delph, 2001).

### Evolvability of stomatal traits

Populations require sufficient genetic variation for selection to act upon for adaptation to occur (Shaw & Etterson, 2001, Jump & Peñuelas, 2005; Cotto et al., 2017). We estimated evolvability for stomatal traits based on the standardization of additive genetic variance and noted that all estimates were close to zero with little to no differences across ecoregions (Table 2). This suggests that populations used in this experiment may have limited capacity to respond to selection. This is a concern in the context of restoration, which will require seed transferred to a new environment to adapt. In *G. triflorum,* limited evolvability in traits associated with water use could lead to adaptational lags when seed is transferred across environments. Reduced evolvability may leave populations more susceptible to demographic declines (Shaw & Etterson, 2001, Jump & Peñuelas, 2005; Cotto et al., 2017). As restorations are multi-species, we advocate for studies that quantify the distribution of genetic variation and evolvability for traits important to adaptation. In this way we may predict long-term evolutionary potential of seed populations used in restoration. Finally, as the work presented here was conducted in a common garden in the prairie region, we urge caution when extrapolating our results into other systems and recognize that a reciprocal transplant experiment allows for the evaluation of additive genetic variance in alvar environments and quantification of plasticity.

## CONCLUSIONS

Identifying how the environment influences the evolution of ecotypes is important to development of seed transfer guidelines. For *G. triflorum* populations, we observed ecoregional differentiation for physiological traits and variation in the distribution of genetic variation. This suggests different seed source populations may exhibit varying evolutionary trajectories that could impact seed transfer decisions. Thus, minimizing environmental differences when transferring seed across environments may be necessary where genetic differences exist among seed sources. However, sourcing local seed may not be enough to create restored populations capable of withstanding climate change (Broadhurst et al., 2008; Bucharova et al., 2018; Espeland et al., 2017). By evaluating heritable genetic variation for traits important to adaptation, it may be possible to quantify the effect selecting seed for restoration beyond local sources will have to long-term adaptive potential.

## ACKNOWLEDGEMENTS

The authors thank Jon Sweetman, Chad Stratilo, Mary Vetter, Rebekah Neufeld, Tyler Stadel, Steve Travers, and the Nature Conservancy of Canada for help with initial field sampling to establish the common garden experiment. In addition, we thank Nick Hugo, Zoe Portlas, Naomi Hegwood, Storm Nies and Zeb Yoko for assistance in the field, and specifically acknowledge Stephen Johnson and Alexis Pearson for help with making stomatal trait impressions in the common garden. This work was supported by a Frank J. Cassel Undergraduate Research Award to K.L.V. and a new faculty award from the office of North Dakota Experimental Program to Stimulate Competitive Research (ND-EPSCoR NSF-IIS-1355466) to J.A.H.

## DATA AVAILABILITY STATEMENT

All data and scripts associated with this manuscript are available on GitHub (https://github.com/KateLVolk/AJB-common-garden-physiology).

## Notes

### Competing Interest Statement

The authors have declared no competing interest.

## REFFERENCES

1. Ahrens, C. W., M. E. Andrew, R. A. Mazanec, K. X. Ruthrof, A. Challis, G. Hardy, M. Byrne, et al. 2020. Plant functional traits differ in adaptability and are predicted to be differentially affected by climate change. Ecology and evolution 10: 232–248.

2. Anderson, J. T., A. M. Panetta, and T. Mitchell-Olds. 2012. Evolutionary and ecological responses to anthropogenic climate change. Plant Physiology 160: 1728–1740.

3. Anderson, J. T., J. H. Willis, and T. Mitchell-Olds. 2011. Evolutionary genetics of plant adaptation. Trends in Genetics 27: 258–266.

4. Anderson, R. H., S. D. Fuhlendorf, and D. M. Engle. 2006. Soil nitrogen availability in tallgrass prairie under the fire–grazing interaction. Rangeland Ecology & Management 59: 625–631.

5. Arntz, M. A., and L. F. Delph. 2001. Pattern and process: evidence for the evolution of photosynthetic traits in natural populations. Oecologia 127: 455–467.

6. Aspinwall, M. J., D. B. Lowry, S. H. Taylor, T. E. Juenger, C. V Hawkes, M. V Johnson, J. R. Kiniry, and P. A. Fay. 2013. Genotypic variation in traits linked to climate and aboveground productivity in a widespread C4 grass: evidence for a functional trait syndrome. New Phytologist 199: 966–980.

7. Baythavong, B. S. 2011. Linking the spatial scale of environmental variation and the evolution of phenotypic plasticity: Selection favors adaptive plasticity in fine-grained environments. American Naturalist 178: 75–87.

8. Becklin, K. M., J. T. Anderson, L. M. Gerhart, S. M. Wadgymar, C. A. Wessinger, and J. K. Ward. 2016. Examining plant physiological responses to climate change through an evolutionary lens. Plant physiology 172: 635–649.

9. Bengtsson, J., J. M. Bullock, B. Egoh, C. Everson, T. Everson, T. O’Connor, P. J. O’Farrell, et al. 2019. Grasslands—more important for ecosystem services than you might think. Ecosphere 10.

10. Bertel, C., P. Schönswetter, B. Frajman, A. Holzinger, and G. Neuner. 2017. Leaf anatomy of two reciprocally non-monophyletic mountain plants (Heliosperma spp.): does heritable adaptation to divergent growing sites accompany the onset of speciation? Protoplasma 254: 1411–1420.

11. Braasch, J. E., L. N. Di Santo, Z. J. Tarble, J. R. Prasifka, and J. A. Hamilton. 2021. Testing for evolutionary change in restoration: A genomic comparison between ex situ, native, and commercial seed sources of Helianthus maximiliani. Evolutionary applications 14: 2206– 2220.

12. Broadhurst, L., M. Driver, L. Guja, T. North, B. Vanzella, G. Fifield, S. Bruce, et al. 2015. Seeding the future - the issues of supply and demand in restoration in Australia. Ecological Management and Restoration 16: 29–32.

13. Bucharova, A., Bossdorf, O., Hölzel, N., Kollmann, J., Prasse, R., & Durka, W. 2019. Mix and match: regional admixture provenancing strikes a balance among different seed-sourcing strategies for ecological restoration. Conservation Genetics 20: 7–17.

14. Bucharova, A., W. Durka, N. Hölzel, J. Kollmann, S. Michalski, and O. Bossdorf. 2017. Are local plants the best for ecosystem restoration? It depends on how you analyze the data. Ecology and Evolution 7: 10683–10689.

15. Catling, Paul M., and V. R. B. 1995. A review of the alvars of the Great Lakes region: distribution, floristic composition, biogeography and protection. Canadian field-naturalist 109: 143–171.

16. Cava, J. A., N. G. Perlut, and S. E. Travis. 2019. Heritability and evolvability of morphological traits of Savannah sparrows (Passerculus sandwichensis) breeding in agricultural grasslands. PloS one 14: e0210472.

17. Chevin, L.-M., R. Lande, and G. M. Mace. 2010. Adaptation, plasticity, and extinction in a changing environment: towards a predictive theory. PLoS biology 8: e1000357.

18. Comer, P. J., J. C. Hak, K. Kindscher, E. Muldavin, and J. Singhurst. 2018. Continent-scale landscape conservation design for temperate grasslands of the Great Plains and Chihuahuan Desert. Natural areas journal 38: 196–211.

19. Cotto, O., J. Wessely, D. Georges, G. Klonner, M. Schmid, S. Dullinger, W. Thuiller, and F. Guillaume. 2017. A dynamic eco-evolutionary model predicts slow response of alpine plants to climate warming. Nature Communications 8: 1–9.

20. Crow, T. M., S. E. Albeke, C. A. Buerkle, and K. M. Hufford. 2018. Provisional methods to guide species specific seed transfer in ecological restoration. Ecosphere 9: e02059.

21. Crowe, K. A., and W. H. Parker. 2008. Using portfolio theory to guide reforestation and restoration under climate change scenarios. Climatic Change 89: 355–370.

22. Didiano, T. J., M. T. J. Johnson, and T. P. Duval. 2016. Disentangling the effects of precipitation amount and frequency on the performance of 14 grassland species. PLoS One 11: e0162310.

23. Dittberner, H., A. Korte, T. Mettler Altmann, A. P. M. Weber, G. Monroe, and J. de Meaux. LJ 2018. Natural variation in stomata size contributes to the local adaptation of water use efficiency in Arabidopsis thaliana. Molecular ecology 27: 4052–4065.

24. Donohue, K. 2009. Completing the cycle: maternal effects as the missing link in plant life histories. Philosophical Transactions of the Royal Society B: Biological Sciences 364: 1059–1074.

25. Drake, P. L., R. H. Froend, and P. J. Franks. 2013. Smaller, faster stomata: scaling of stomatal size, rate of response, and stomatal conductance. Journal of experimental botany 64: 495– 505.

26. Dudley, S. A. 1996. Differing selection on plant physiological traits in response to environmental water availability: a test of adaptive hypotheses. Evolution 50: 92–102.

27. Espeland, E. K., N. C. Emery, K. L. Mercer, S. A. Woolbright, K. M. Kettenring, P. Gepts, and J. R. Etterson. 2017. Evolution of plant materials for ecological restoration: insights from the applied and basic literature. Journal of Applied Ecology 54: 102–115.

28. Etterson, J. R., and R. G. Shaw. 2001. Constraint to adaptive evolution in response to global warming. science 294: 151–154.

29. Falconer, D. S. 1996. Introduction to quantitative genetics. Pearson Education India.

30. Galliart, M., N. Bello, M. Knapp, J. Poland, P. St Amand, S. Baer, B. Maricle, et al. 2019. Local adaptation, genetic divergence, and experimental selection in a foundation grass across the US Great Plains’ climate gradient. Global Change Biology 25: 850–868.

31. Gascoigne, W. R., D. Hoag, L. Koontz, B. A. Tangen, T. L. Shaffer, and R. A. Gleason. 2011. Valuing ecosystem and economic services across land-use scenarios in the Prairie Pothole Region of the Dakotas, USA. Ecological Economics 70: 1715–1725.

32. Ghalambor, C. K., J. K. McKay, S. P. Carroll, and D. N. Reznick. 2007. Adaptive versus non adaptive phenotypic plasticity and the potential for contemporary adaptation in new environments. Functional ecology 21: 394–407.

33. Gilbert, A. L., and D. B. Miles. 2019. Antagonistic responses of exposure to sublethal temperatures: adaptive phenotypic plasticity coincides with a reduction in organismal performance. The American Naturalist 194: 344–355.

34. Guo, C., Ma, L., Yuan, S., & Wang, R. 2017. Morphological, physiological and anatomical traits of plant functional types in temperate grasslands along a large-scale aridity gradient in northeastern China. Scientific reports 7: 1–10.

35. Hamilton, J. A., and C. G. Eckert. 2007. Population genetic consequences of geographic disjunction: A prairie plant isolated on Great Lakes alvars. Molecular Ecology 16: 1649– 1660.

36. Hamilton, J., S. Flint, J. Lindstrom, K. Volk, R. Shaw, and M. Ahlering. 2020. Evolutionary approaches to seed sourcing for grassland restorations. New Phytologist 225: 2246–2248.

37. Hansen, T. F., and D. Houle. 2008. Measuring and comparing evolvability and constraint in multivariate characters. Journal of evolutionary biology 21: 1201–1219.

38. Hansen, T. F., C. Pélabon, and D. Houle. 2011. Heritability is not evolvability. Evolutionary Biology 38: 258–277.

39. Havens, K., P. Vitt, S. Still, A. T. Kramer, J. B. Fant, and K. Schatz. 2015. Seed sourcing for restoration in an era of climate change. Natural Areas Journal 35: 122–133.

40. Hoekstra, J. M., T. M. Boucher, T. H. Ricketts, and C. Roberts. 2005. Confronting a biome crisis: global disparities of habitat loss and protection. Ecology letters 8: 23–29.

41. Jump, A. S., and J. Penuelas. 2005. Running to stand still: adaptation and the response of plants to rapid climate change. Ecology letters 8: 1010–1020.

42. Kawecki, T. J., & Ebert, D. 2004. Conceptual issues in local adaptation. Ecology letters 7: 1225– 1241.

43. Kittelson, P. M., S. Wagenius, R. Nielsen, S. Qazi, M. Howe, G. Kiefer, R. G. Shaw, and N. J. B. Kraft. 2015. How functional traits, herbivory, and genetic diversity interact in Echinacea: Implications for fragmented populations. Ecology 96: 1877–1886.

44. Kulbaba, M. W., S. N. Sheth, R. E. Pain, V. M. Eckhart, and R. G. Shaw. 2019. Additive genetic variance for lifetime fitness and the capacity for adaptation in an annual plant. Evolution 73: 1746–1758.

45. Kulbaba, M. W., Z. Yoko, and J. A. Hamilton. 2021. Chasing the fitness optimum: temporal variation in the genetic and environmental expression of life-history traits for a perennial plant. *bioRxiv*.

46. Lande, R. 1992. Neutral theory of quantitative genetic variance in an island model with local extinction and colonization. Evolution 46: 381–389.

47. Levins, R. 1963. Theory of fitness in a heterogeneous environment. II. Developmental flexibility and niche selection. The American Naturalist 97: 75–90.

48. Lowry, D. B., M. C. Hall, D. E. Salt, and J. H. Willis. 2009. Genetic and physiological basis of adaptive salt tolerance divergence between coastal and inland Mimulus guttatus. New Phytologist 183: 776–788.

49. March-Salas, M., M. van Kleunen, and P. S. Fitze. 2019. Rapid and positive responses of plants to lower precipitation predictability. Proceedings of the Royal Society B 286: 20191486.

50. Maricle, B. R., K. L. Caudle, K. J. Lindsey, S. G. Baer, and L. C. Johnson. 2017. Effects of extreme drought on photosynthesis and water potential of Andropogon gerardii (big bluestem) ecotypes in common gardens across Kansas. Transactions of the Kansas Academy of Science 120: 1–16.

51. McKay, J. K., C. E. Christian, S. Harrison, and K. J. Rice. 2005. “How local is local?”—a review of practical and conceptual issues in the genetics of restoration. Restoration Ecology 13: 432–440.

52. McMillan, C. 1959. The role of ecotypic variation in the distribution of the central grassland of North America. Ecological Monographs 29: 286–308.

53. Olsen, J. T., K. L. Caudle, L. C. Johnson, S. G. Baer, and B. R. Maricle. 2013. Environmental and genetic variation in leaf anatomy among populations of Andropogon gerardii (Poaceae) along a precipitation gradient. American Journal of Botany 100: 1957–1968.

55. Picotte, J. J., D. M. Rosenthal, J. M. Rhode, and M. B. Cruzan. 2007. Plastic responses to temporal variation in moisture availability: consequences for water use efficiency and plant performance. Oecologia 153: 821–832.

56. Ramírez Valiente, J. A., N. J. Deacon, J. Etterson, A. Center, J. P. Sparks, K. L. Sparks, T. LJ Longwell, et al. 2018. Natural selection and neutral evolutionary processes contribute to genetic divergence in leaf traits across a precipitation gradient in the tropical oak Quercus oleoides. Molecular Ecology 27: 2176–2192.

57. Reed, T. E., R. S. Waples, D. E. Schindler, J. J. Hard, and M. T. Kinnison. 2010. Phenotypic plasticity and population viability: the importance of environmental predictability. Proceedings of the Royal Society B: Biological Sciences 277: 3391–3400.

58. Singmann, H., Bolker, B., Westfall, J., Aust, F., Ben-Shachar, M.S., Hojsgaard, S., Fox, J., Lawrence, M.A., Mertens, U., Love, J., Lenth, R. , Christensen, R. H. B. 2021. Package ‘afex’.

59. VanWallendael, A., D. B. Lowry, and J. A. Hamilton. 2022. One hundred years into the study of ecotypes, new advances are being made through large-scale field experiments in perennial plant systems. Current Opinion in Plant Biology 66: 102152.

60. Wang, T., Hamann, A., Spittlehouse, D., & Carroll, C. 2016. Wang T, Hamann A, Spittlehouse D, Carroll C. Locally downscaled and spatially customizable climate data for historical and future periods for North America. PloS one 11.

61. Wang, T., G. A. O’Neill, and S. N. Aitken. 2010. Integrating environmental and genetic effects to predict responses of tree populations to climate. Ecological applications 20: 153–163.

62. Wimberly, M. C., D. M. Narem, P. J. Bauman, B. T. Carlson, and M. A. Ahlering. 2018. Grassland connectivity in fragmented agricultural landscapes of the north-central United States. Biological Conservation 217: 121–130.

63. Yoko, Z. G., K. L. Volk, N. A. Dochtermann, and J. A. Hamilton. 2020. The importance of quantitative trait differentiation in restoration: landscape heterogeneity and functional traits inform seed transfer guidelines. AoB Plants 12: plaa009.

64. Young, A., T. Boyle, and T. Brown. 1996. The population genetic consequences of habitat fragmentation for plants. Trends in ecology & evolution 11: 413–418.

